# dtangle: accurate and fast cell-type deconvolution

**DOI:** 10.1101/290262

**Authors:** Gregory J. Hunt, Saskia Freytag, Melanie Bahlo, Johann A. Gagnon-Bartsch

## Abstract

**Motivation:** Understanding cell type composition is important to understanding many biological processes. Furthermore, in gene expression studies cell type composition can confound differential expression analysis (DEA). To aid understanding cell type composition, methods of estimating (deconvolving) cell type proportions from gene expression data have been developed.

**Results:** We propose dtangle, a new cell-type deconvolution method. dtangle works on a range of DNA microarray and bulk RNA-seq platforms. It estimates cell-type proportions using publicly available, often cross-platform, reference data. To comprehensively evaluate dtangle, we assemble ten benchmark data sets. Here, dtangle is competitive with published deconvolution methods, is robust to selection of tuning parameters and is quicker than other methods. As a case study, we investigate the human immune response to Lyme disease. dtangle’s estimates reveal a temporal trend consistent with previous findings and are important covariates for DEA across disease status.

**Availability:** dtangle is on CRAN (cran.r-project.org/package=dtangle) or github (dtangle.github.io).

## 1 Introduction

Complex organisms have a vast collection of specialized cell types. The presence and interaction of these cell types is important to understanding many biological processes. For example, shifts in the relative composition of cell types is important to developmental processes of organisms including embryogenesis, morphogensis, cell differentiation and growth (Lu *et al.*, 2003). Likewise, understanding the presence or absence of cell types is of direct etiological interest for many diseases and dysfunctions (Newman *et al.*, 2015; Abbas *et al.*, 2009; Altboum *et al.*, 2014; Lu *et al.*, 2003). For example, changes in glial populations in brain tissue are characteristic of Alzheimer’s disease (Mohammadi *et al.*, 2015). Similarly, white blood cell composition can be indicative of acute cellular rejection of transplanted kidneys (Shen-Orr *et al.*, 2010). Cell type composition is also important in tumorigenic processes. It has been shown that heterogeneity of tumors cells is implicated in the metastatic potential of cancer (Marusyk and Polyak, 2011; Lu *et al.*, 2003).

Given the importance of understanding cell type composition, several methods to estimate cell type proportions using high-throughput gene profiling experiments have been developed. Known as “cell type deconvolution”, these methods have been successfully employed in a variety of applications. Deconvolution algorithms have been used to study cell type compositional changes in patients in clinical studies (Newman *et al.*, 2015; Abbas *et al.*, 2009; Gong *et al.*, 2011; Altboum *et al.*, 2014; Bowling *et al.*, 2017). In these studies, estimating constituent cell types of carefully selected tissues reveals important cell type compositional dynamics of diseases. Similarly, such gene expression deconvolution has been posited as useful for clinical cell type monitoring (Newman *et al.*, 2015). Tracking leukocytes, for example, is known to be a useful clinical tool and something of which deconvolution algorithms are capable. Finally, estimating cell type proportions is important for deconfounding differential expression analysis. In differential expression studies detecting gene expression differences within each cell type is confounded by changes in the cell type composition across the factor of interest. For example, diseases will simultaneously affect changes in gene expression within each cell type and through compositional changes in the tissues. Including estimated proportions of cell types to account for this confounding has been shown to improve differential expression analysis (Capurro *et al.*, 2015; Hagenauer *et al.*, 2016).

We present dtangle, a new deconvolution method that is accurate, robust, and simple to compute. It estimates cell-type proportions using biologically plausible models of high throughput profiling technology. We comprehensively compare dtangle to other methods on 10 benchmark data sets. These data sets include many different cell types, profiling technologies, and cover realistic scenarios like batch effects, mixed technologies, and third party references. Analysis of this data shows that dtangle out-competes existing methods in a broad range of applications.

## 2 Materials and Methods

dtangle assumes a linear mixing process of actual gene expressions (mRNA transcripts) in each sample and combines this with a linear model connecting the actual gene expressions to the measured gene expressions from the profiling technology. dtangle combines these models with a precise definition of marker genes.

dtangle requires two pieces of external knowledge: (1) reference data and (2) marker genes. First, dtangle requires auxiliary gene expression reference data for each cell type presumed to be in the mixture samples. In realistic use cases this auxiliary reference data can come from online repositories of gene expression data like GEO (Edgar, 2002). Second, dtangle requires marker genes. A marker gene is a gene which is expressed predominantly by only one cell type. If desired, these marker genes are automatically determined by dtangle. The user need only specify the number of markers genes to use as a tuning parameter. Otherwise, the marker genes may be specified explicitly by the user.

### 2.1 The dtangle Estimator

Assume we have a mixture sample of *K* cells types. Let *Y* ∈ ℝ^*N*^ be the (base-2) log-scale expression measurements of this mixture sample and *p*_1_, *…, p*_*K*_ be the mixing proportions of the cell-types. For *k* = 1, *…, K* assume that there are *ν*_*k*_ reference samples of cell type *k* and let *Z*_*kr*_ ∈ ℝ^*N*^ be the log-scale expressions of the *r*^*th*^ type *k* reference. Furthermore, let *G*_*k*_ ⊂ {1, *…, N*} be the set of type *k* marker genes. These marker gene sets are mutually disjoint.

For *g*_*k*_ = *|G*_*k*_*|* define 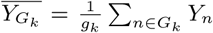 and 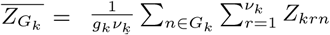 to be the average of all type *k* marker genes across the mixture and reference samples, respectively. Given a technological sensitivity parameter *γ* ∈ ℝ^+^ define

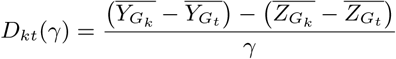

and *D*_*k*_(*γ*) = (*D*_*k*1_(*γ*), *…, D*_*kK*_(*γ*)). The elements *D*_*kt*_(*γ*) are a normalized measure of the type *k* marker genes’ expression over the type *t* markers’ expressions in the mixture. Precisely, it is the difference of marker expressions, 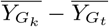, baseline normalized by their difference across the references, 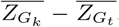, and the technological sensitivity, *γ*. Since the (normalized) relative expressions of marker genes in the mixture *D*_*k*_(*γ*) relates to the relative abundance of cell types, we estimate of *p*_*k*_ by mapping *D*_*k*_(*γ*) ∈ ℝ into the unit interval [0, 1] by a multivariate logistic function *L*_*k*_ : ℝ^*K*^ *→* [0, 1]. Precisely, for *x* ∈ ℝ^*K*^ let

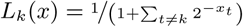

then we estimate *p*_*k*_ as

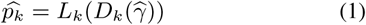

for some estimator 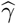 of *γ* (see Supplement for details of estimating *γ*). This definition ensures that 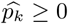 and 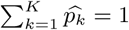.

### 2.2 Terminology, Transformations and Scales

To relate dtangle’s approach to that of existing methods we first define some terminology. Figure 1 guides this discussion. Measured expressions are assigned by a gene expression profiling technology (GEP) by measuring the amount of mRNA transcribed from each gene. Typically these measured expressions are further summarized and transformed by algorithms like MAS5.0 or RMA. These processed measurements are called the “measured gene expressions.” Often, they are transformed by a logarithm to produce “log scale” measure expressions, otherwise, they are “linear scale”. We call the true, yet unobserved, amount of mRNA transcribed from each gene the “actual expression” of the gene. This actual gene expression can be considered on the linear scale or the log scale.

**Figure 1:**
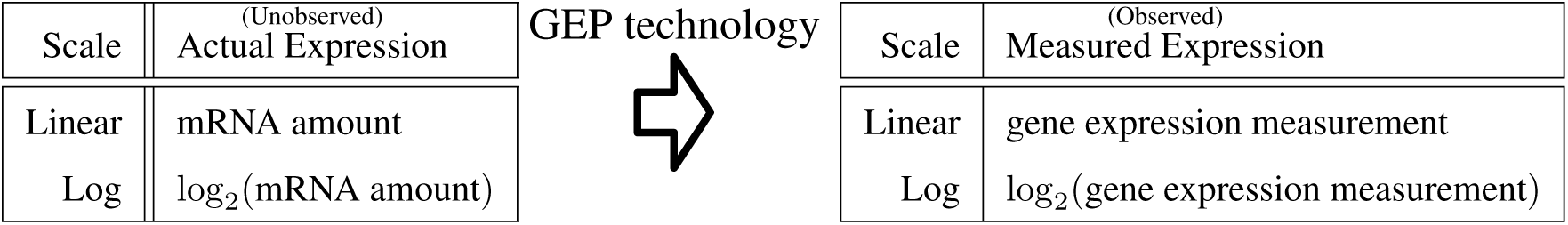
Measured expressions (log or linear) arise from a measurement process on the actual expressions (log or linear).

### 2.3 dtangle’s Statistical Model

dtangle’s approach combines three components:

1. dtangle assumes that, on the linear scale, the actual expressions in the mixture sample are a linear mixture of the actual expressions in the reference samples.
2. dtangle linearly models the GEP measurement process on the log scale so that the log measured expressions arise linearly from the log actual expressions.
3. dtangle precisely defines marker genes.

We consider each of these components in turn.

dtangle’s first component is a linear mixing model of actual expressions. If *η*_*kn*_ is the actual linear-scale expression of the *n*^*th*^ gene in a sample of type *k* cells then we assume

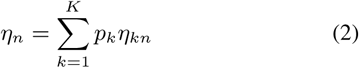

where *ηn* is the linear-scale actual expression of gene *n* in the mixture. That is, the total amount of a transcript in the mixture is the sum of the amount of that transcript from each cell type.

Second, dtangle models how measured expressions arise from actual expressions. Since the *Y′s* and *Z′s* are on the log scale and the *η′s* are not, we formulate this statistically as

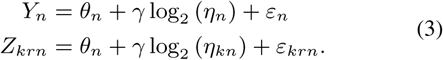

for *n* = 1, *…, N, r* = 1, *…, ν*_*k*_ and *k* = 1, *…, K*. The intercept *θ*_*n*_ depends on the gene, the slope (sensitivity parameter) *γ* is universal to all genes. The *ε* error terms are uncorrelated with mean zero and finite variance.

Finally dtangle defines marker genes so that if *n* is a marker gene for cell type *k* (*n* ∈ *G*_*k*_), then

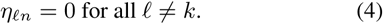

That is, if *n* is a marker gene of type *k* then its actual expression is zero in any cell type except type *k*. (We don’t expect this to hold exactly in reality, but at least approximately.)

Combining Equation (2) with Equation (3) and Equation (4) we have

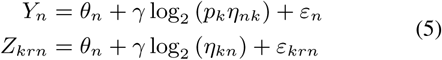

for all *n* ∈ *G*_*k*_. Define

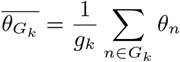

Then

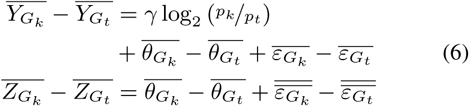

Where 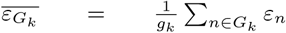 and 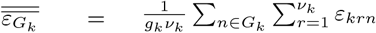. Further let

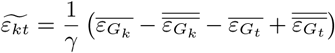

and note that 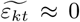 when the number of marker genes is large. Now we have that

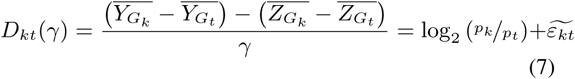

All together then

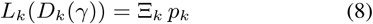

where 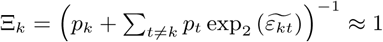.

The dtangle estimate 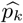 is the plug-in estimator replacing *γ* with 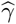 in Equation (8), 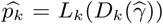. Thus to the extent 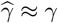 and Ξ_*k*_ *≈* 1 then 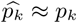.

### 2.4 Relationship of dtangle to other deconvolution methods

Existing deconvolution methods assume a linear mixing model on the measured expressions. The model is

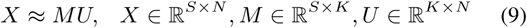

where *X* is the mixture samples’ expressions, *M*_*sk*_ is the percentage of type *k* cells in sample *s*, and the *K* rows of *U* are the typical expressions of the *K* cell types.

Given *X*, the deconvolution problem is estimating either (a) the mixing proportions *M*, if *U* is known, (b) *U*, if *M* is known, or (c) *U* and *M* jointly, if neither are known. All three problems are considered instances of deconvolution. dtangle most closely resembles problem (a), of estimating *M* given *X* and *U*. In the Results section we compare dtangle to methods solving both (a) and (c) since they also estimate *M*. Problem (a), called “partial deconvolution”, is typically solved as a regression or penalized regression problem (Abbas *et al.*, 2009; Gong *et al.*, 2011; Lu *et al.*, 2003; Wang *et al.*, 2006; Qiao *et al.*, 2012; Altboum *et al.*, 2014; Newman *et al.*, 2015), problem (c), called “full deconvolution”, is usually accomplished by non-negative matrix factorization (Venet *et al.*, 2001; Repsilber *et al.*, 2010; Gaujoux and Seoighe, 2012; Zhong *et al.*, 2013).

#### 2.4.1 Choice of Scale: Interpretability, Robustness, and Efficiency

Existing partial deconvolution methods fit Equation (9) by some form of linear regression on either the linear or log scale. We assert that a linear model on linear measured expressions is physically plausible. The model assumes that mRNA from a sample of cells is the sum of the mRNA from each cell. While the model is plausible, fitting the model by linear regression on the linear scale is not robust and is statistically inefficient. The linear expressions are very right-skewed and so the fit is dominated by the tail (Li *et al.*, 2016). Furthermore, heteroskedasticity means regression is sub-optimal as variance of expressions typically scales with the mean (Li *et al.*, 2016). Fitting Equation (9) on log expressions is more robust since the transformation ameliorates the skewness and heteroskedasticity. However, critically, Equation (9) is not physically meaningful on a log scale. It is tantamount to saying that the mRNA in a mixture sample is the product (not sum) of the mRNA from each cell. We assert that this does not make sense physically.

dtangle’s approach is to take advantage of the beneficial aspects of each scale while avoiding their problems. First, dtangle’s linear mixing model of linear-scale actual expressions is biologically interpretable. Second, dtangle’s linear model connecting actual and measured expression on the log scale is robust and efficient. Working on this scale is robust because we average before exponentiating.

## 3 Results

### 3.1 Benchmarking

In order to evaluate dtangle we compare it to seven other deconvolution algorithms (Supplementary Table S1). To ensure an accurate comparison we compare to six methods with a standardized interface from the well-tested CellMix package in R (Gaujoux, 2013). We also compare to CIBERSORT because it is a recent and powerful method (Newman *et al.*, 2015). Note that we only compare dtangle against methods that estimate cell-type proportions from gene expression data for arbitrary cell types. We do not compare to methods like xCell (Aran *et al.*, 2017) which produce enrichment scores and not percentages. We also do not compare against the many deconvolution methods for methylation data or fully-unsupervised methods whose cell types have to be inferred with further post-hoc analysis e.g. CAM (Wang *et al.*, 2016). Furthermore, we do not compare against methods that only estimate cell-type proportions from a very specific subset of cells or only in the context of a specific problem, for example, immune cell infiltration of tumors by methods like TIMER (Li *et al.*, 2016).

Like dtangle, all methods we survey require marker genes. However four methods require only marker genes and do not explicitly require reference data on each cell type. Following Gaujoux (2013) we call the latter methods “full” deconvolution and the former “partial” deconvolution (Supplementary Table S1). While the full deconvolution methods do not explicitly require references, marker genes are typically found through DEA on reference data. Furthermore, there has been some development of “completely unsupervised” deconvolution methods (e.g. Wang *et al.* (2016)) that claim to require neither markers nor references. However the results from these algorithms are uninterpretable biologically unless references are used post-hoc to map proportions to cell types. For this reason we do not compare to such methods. In any case, *de facto* all methods (full, partial or fully-unsupervised) use reference data in one way or another.

While the full deconvolution algorithms also estimate average type-specific gene expressions we only compare dtangle to their estimated mixing proportions, since this is all that dtangle estimates. We choose marker genes for deconvolution following Abbas *et al.* (2009) This ranks genes by the *p*-value from a *t*-test between the two most highly expressed cell types for each gene. We then designate the top 10% of genes for each cell type as markers. The same markers are used for each deconvolution algorithm.

### 3.2 Data Sets Compared

We compare dtangle to seven algorithms across ten benchmarking data sets (Supplementary Table S2). This is the largest ever meta analysis (in both number of algorithms and number of data sets) of partial deconvolution algorithm accuracy. Furthermore, it is the first meta analysis to evaluate methods on both microarray and RNA-seq platforms. For each data set the true mixing proportions are known, either because the experiment was conducted by mixing each cell type in the reported proportions or because an independent physical sorting technique, like flow cytometry, was used to estimate the cell type mixing proportions. The majority of data sets include their own reference samples.

For the microarray experiments we base-2 logarithmically transform the expression measurements. For the RNA-seq data we analyze the base-2 logarithm of the read count plus one. All data was quantile normalized. Detailed information and processing source code is available for each data set as part of our dtangle.data R package available at dtangle.github.io.

### 3.3 Mixture Experiments With References

#### 3.3.1 Microarray

We consider data sets Abbas, Kuhn, Gong, Shi and Shen-Orr (Supplementary Table S2) that are microarray mixture experiments. For each algorithm we estimate the mixing proportions in each data set and compute the mean of the absolute value of the error of each proportion from the truth. dtangle has the lowest mean and median error across the data sets (Figure 2a). Furthermore dtangle has the lowest spread. This meta-analysis shows that for the microarray mixture experiments dtangle is not only the most accurate algorithm but it is also the most consistently accurate algorithm.

**Figure 2:**
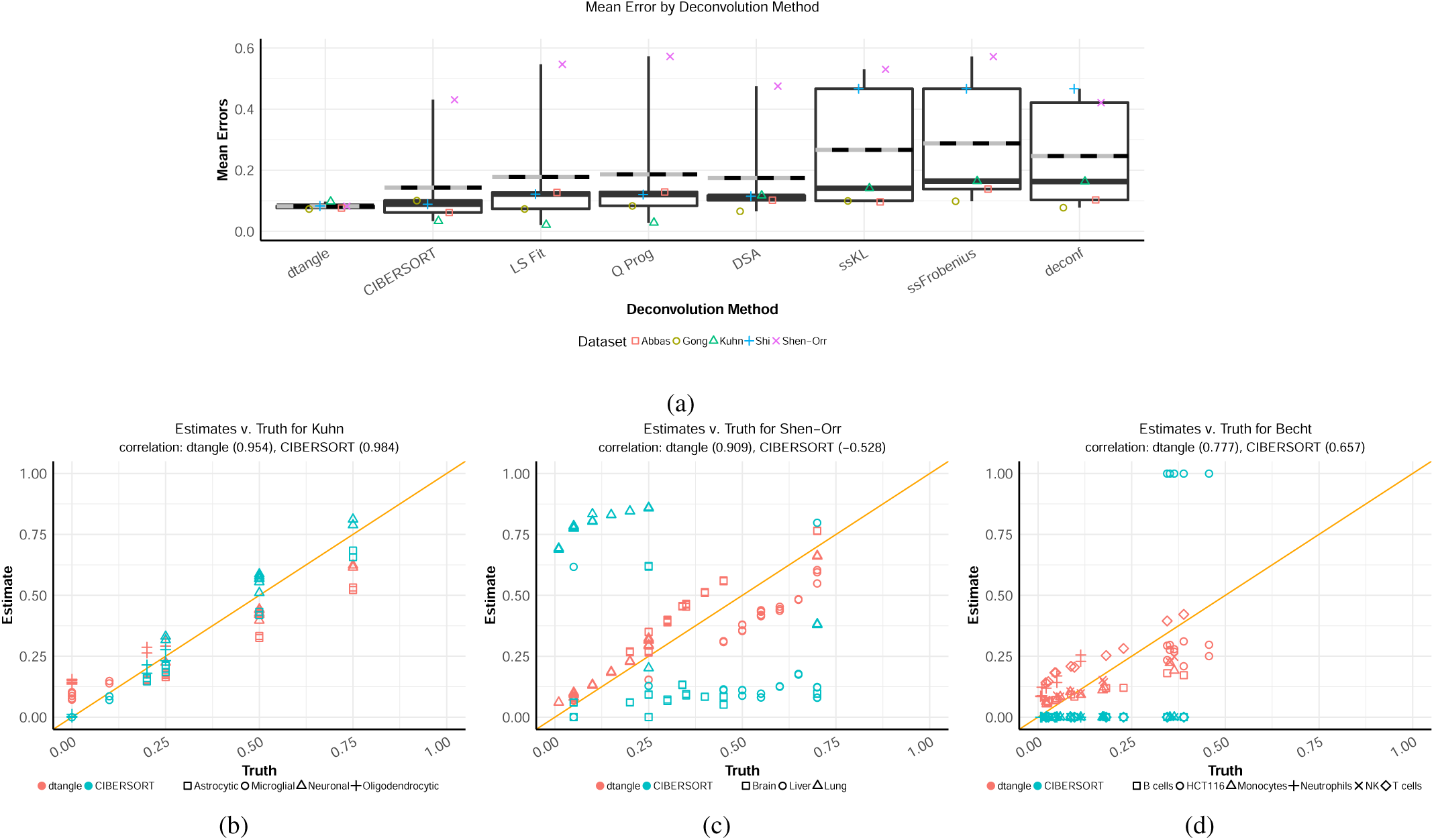
(a) Side-by-side box plots of the mean absolute errors across the algorithms. The bold line is median, striped line is mean. Overlapping are jittered points of the mean error for each data set. (b-c) Scatter plots of dtangle and CIBERSORT. Each point is a are cell type in a sample.

We use scatter plots to compare dtangle and CIBERSORT on two data sets where dtangle performs worst and best relative to other algorithms (Figure 2b and Figure 2c). We compare to CIBERSORT as it is the current state-of-the-art and the second best performing algorithm after dtangle. For the Kuhn data four types of rat brain cells (astrocytic, microglial, neuronal, and oligodendrocytic) were mixed in known proportions. While dtangle does poorly relative to other deconvolution algorithms it still performs quite well. The estimated mixing proportions are still highly correlated with the truth (*r* = 0.954).

The Shen-Orr data is from a microarray mixture experiment where rat liver, brain and lung cDNA were mixed in known proportions. dtangle strongly outperforms all other algorithms. Even a well performing algorithm like CIBERSORT performs relatively poorly on this microarray mixture experiment (Figure 2c). In particular, CIBERSORT seems to struggle distinguishing liver and lung cell signatures.

#### 3.3.2 RNA-seq

We also investigate the performance of deconvolution methods on RNA-seq mixture experiments (Supplementary Figure S2). There is no clear best algorithm across the data sets, however the partial deconvolution algorithms (including dtangle) typically outperform the full deconvolution methods.

### 3.4 Mixtures Without References

In practice pure reference samples of each cell type are not typically generated along with the mixed samples to be deconvolved. In this case existing reference data for each of the cell types to deconvolved must be procured. Typically these pure reference samples are collected from repositories like GEO.

The Becht data set is a mixture experiment where known quantities of cDNA from the HCT116 colorectal carcinoma line and various leukocytes (NK, B, neutrophils, T, and monocytes) were mixed in known quantities and analyzed with an Affymetrix microarray. Unlike previous data sets no reference data was produced as part of the mixture experiment. Like the authors we use publicly available expression data from GEO as references for each cell type. In total there are 776 samples gathered from GEO which we use to create reference profiles for the six cell types.

dtangle performs much better than the other algorithms on this data set (Supplementary Figure S1). The estimated proportions are much more highly correlated with the truth when using dtangle (*r* = 0.777) than CIBERSORT (*r* = 0.657) (Figure 2d).

### 3.5 Performance evaluation with flow cytometry based cell sorting

Mixture experiments are only a surrogate for cell mixtures found in organisms. Realistically, deconvolution methodology is applied to complex tissue extracted from an organism. Such tissue will be a mixture of many cell types (more types than in a typical mixture experiment) and the cell types will have complex inter-cellular interactions modifying their gene expressions. The difficulties in estimating cell type proportions from such complex tissue is likely only partially explored by a mixture experiment.

The Newman follicular lymphoma (FL) data was generated by taking lymph node biopsy samples and enumerating immune cell sub-types using flow cytometry (Newman *et al.*, 2015). This process identified 3 leukocyte types (B, CD4 T and CD8 T) in various proportions across samples from 14 patients. As cell type expression reference data we use the LM22 reference data set compiled by Newman *et al.* (2015). It contains gene expressions of 22 white blood cell types as references. Similar to Newman *et al.* (2015) we group these 22 types into 12.

The Newman peripheral blood mononuclear cells (PBMC) data was generated from blood samples from twenty adults where the proportions of nine types of leukocytes were determined by flow cytometry. We again use the LM22 data set Newman *et al.* (2015) as reference profiles for these cell types. dtangle compares well with other deconvolution methods on these two Newman data sets (Supplementary Figure S7a, S7b)

The Newman FL and Newman PBMC data sets are our most realistic applications of dtangle and the other deconvolution algorithms. Not surprisingly the accuracy of the deconvolution algorithms is lower on these realistic data sets than most of the mixture experiments. We see some high outliers for the Newman FL data set. There are several factors that make these data sets difficult. Firstly since the data sets are not mixture experiments, and hence the true mixing proportions are unknown, we compare each algorithm’s performance to the percentages reported by flow cytometry, which has its own sources of error (Saeys *et al.*, 2016). Secondly, an additional factor that makes these data sets difficult to correctly deconvolve is the number and close relationship of the cell types. In both experiments we are deconvolving many cell types including some whose presence was not reported by flow cytometry. Trying to deconvolve many cell types is more difficult than deconvolving only a few cell types. If we restrict the cell types we search for to only those cell types that are reported by flow cytometry then accuracy generally increases. For example, the B cells may be more accurately estimated in the Newman FL data (Supplementary Figure S3 -S4). This suggest that, in practice, researchers may be able to improve accuracy by narrowing down the cell types assumed to comprise the sample. For example, we know that certain cell types will be filtered out in PBMC samples and so accuracy could be improved by removing them from consideration.

A final feature of these realistic data sets is that they use external reference profiles for the cell types. Since these external references are not created by the same experiment there are often large batch effects one needs to account for. In these experiments we quantile normalize the data to account for such batch effects. Unfortunately quantile normalization is a nonlinear transformation that does not always work well with the deconvolution algorithms. In particular, this quantile normalization worsens the large outliers of the B cells for the FL data (Supplementary Figure S7a). Quantile normalizing increases the estimating accuracy for some cell types (e.g. gamma-delta T) but decreases it for others (e.g. CD4 T) (Supplementary Figure S5 -S6). The median error decreases if we do not quantile normalize, however some sort of normalization is clearly needed. This illustrates the need for further investigation of the best normalization methods for deconvolution applications, a topic for which there is little existing literature.

### 3.6 Meta-analysis

We compare dtangle to the other algorithms in a meta-analysis (Figure 3). dtangle has the lowest mean and median error of all methods. It also has the lowest maximum error and is consistently accurate across all data sets. dtangle is a general purpose cell type deconvolution algorithm that works well across many technologies and tissue types.

**Figure 3:**
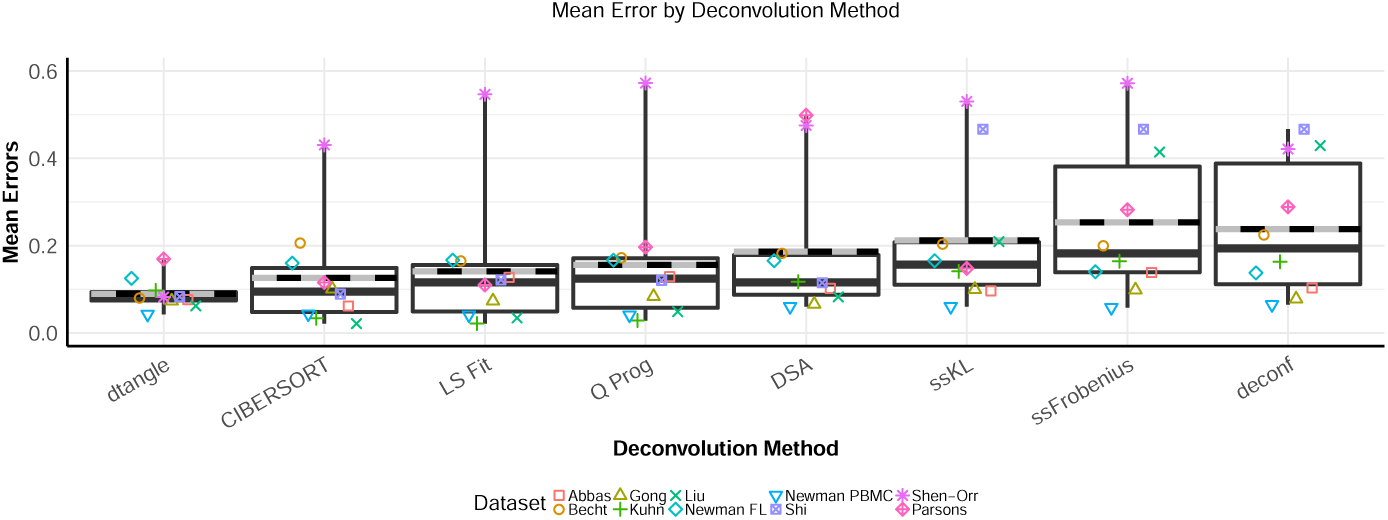
Meta-analysis of deconvolution algorithms. Side-by-side box plots of the mean errors across the algorithms. The bold line is median, striped line is mean. Overlapping are jittered points of the mean error for each data set.

### 3.7 Robustness to marker selection

Thus far we have been following Abbas *et al.* (2009), ranking marker genes with a t-test p-value between the top two most expressed cell types for each gene. To analyze the sensitivity of dtangle to how the markers are ranked we consider another way of ranking marker genes. This second method looks, for each gene, at the ratio of the mean expression for each cell type to the sum of the mean expressions of the gene by all other cell types. We call the Abbas method “p-value” and call this latter approach “Ratio.”

In Figure 4 we look at the grand error mean of each algorithm across all data sets for a range of marker tuning parameters. We compare partial deconvolution algorithms as they are the most competitive with dtangle. dtangle is robust to the way markers are ranked (p-value or Ratio) and ranking threshold (quantile cutoff) determining the number of markers to use. CIBER-SORT appears to have a strong dependence on the quantile cutoff especially when using p-value ranking. The grand error means of LS Fit and Q Prog are sensitive to which ranking method is chosen (they are better using p-value).

**Figure 4:**
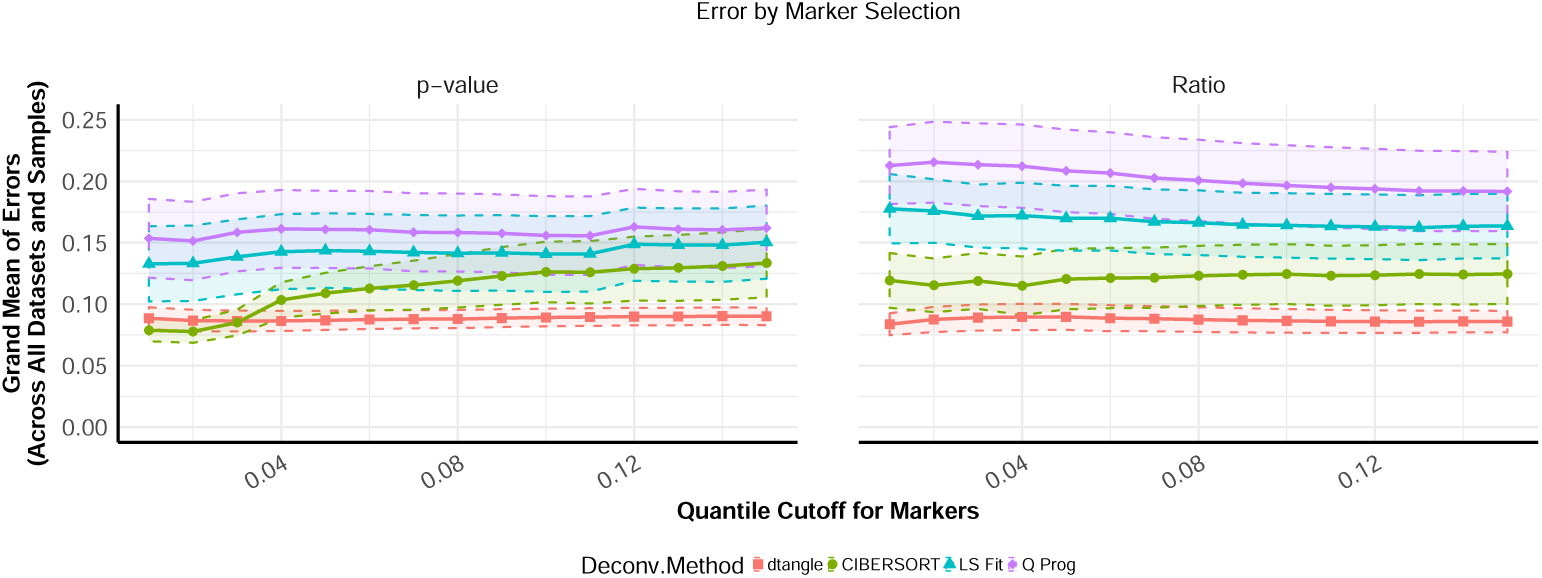
Grand error means across marker ranking methods (p-value and Ratio) and number of markers. Approximate 80% confidence bands included.

Marker gene selection also influences computational time. For each data set we timed all algorithms across a range of quantile cutoffs using *p*-value ranking (Supplementary Figure S8). dtangle is consistently the fastest algorithm. It is between one and four orders of magnitude faster than other algorithms regardless of what quantile cutoff is used. The slowest algorithms are iterative approaches like CIBERSORT, ssKL, ssFrobenius, DSA and deconf. Depending on the number of markers, these iterative methods may take from half-an-hour up to an hour or more to converge on a 4.00 GHz processor. dtangle never takes more than about 10^*-*2^ seconds. Speed is important to practitioner workflow and easily reproducible research. This is especially true for re-analysis of large historical data collections. For example, deconvolution of the more than 70,000 data sets comprising recount2 will necessitate a highly efficient method (Collado-Torres *et al.*, 2017).

### 3.8 Application To Lyme Disease

To demonstrate dtangle on a biological problem we consider RNA-seq data of PBMCs from Lyme disease patients (Bouquet *et al.*, 2016). To better understand persistent Lyme symptoms (e.g. fatigue or arthritis) it is of interest to understand the progression of the human immune response to Lyme (Bouquet *et al.*, 2016). To this end Bouquet *et al.* measure gene expression in a subset of white blood cells (PBMCs). PBMCs of 28 patients were collected at the point of diagnosis (V1), after a 3-week course of doxycycline (V2) and 6 months later (V5). PBMCs from 13 matched controls were also collected (C).

We use dtangle to estimate, for each sample, the celltype proportions of nine types of PBMCs (B, dendritic, macrophages, mast, monocytes, NK, CD4 T, CD8 T and gamma-delta T). We use as reference the LM22 data set from Newman *et al.* (2015), choosing the top 10% of differentially expressed genes for each cell type as markers. We find that the phagocytes (dendritic, macrophages, mast and monocytes) make up a larger percentage of the patients’ PBMCs earlier, rather than later, in the infection (Supplementary Figure S9). We see a large difference between the control group and V1 and decreasing differences between the controls and V2 and V5. Natural killer (NK) cells follow this same pattern.

The estimated cell type percentages agree with the current understanding of Lyme. The initial infection induces an immune response where fast-acting phagocytes are recruited to attack the foreign bacteria (Dame *et al.*, 2007). This agrees with dtangle’s estimates of a relatively large percentage phagocytes early in the infection that decreases with time. Phagocytes decrease in numbers once the bacteria has been cleared and they are no longer needed. Furthermore, NK cells follow the same pattern. This agrees with work from Horowitz *et al.* (2012) showing NK cells are rapidly activated by cytokines after a bacterial infection.

In Bouquet *et al.* (2016) the authors seek to find genes that are differentially expressed among the groups (V1, V2, V5 and C). Following Bouquet *et al.* (2016) we compare the control group to V1, V2 and V5 and find that there are 399 genes that are differentially expressed in the intersection of each of the three comparisons. This was done controlling for a FDR of 0.05 by the Benjamini-Hochberg procedure.

As this previous differential expression analysis was not corrected for cell type proportions we expect to find genes that are correlated with cell type. We add in covariates to account for composition of fast-acting cell types (phagocyte and NK). After doing so we only find 158 genes differentially expressed in the same comparison. Thus the cell type composition changes the results of the analysis greatly. dtangle is one tool practitioners can use to help tease apart histological changes in cell composition from changes in gene expression within particular cell types.

## 4 Discussion

dtangle is a simple and robust deconvolution estimator. It is a closed-form estimator deriving from plausible biological modeling. Our meta-analyses show that dtangle is a robust and accurate, typically performing better than seven of the best existing methods across ten diverse data sets. It can accurately deconvolve cell types using microarray and RNA-seq technology and is very fast to compute where other methods are not. Furthermore it is consistent with standard physical sorting methods like flow cytometry on realistic complex clinical tissue. Finally, dtangle has competitive accuracy when dealing with realistic data sets where the reference samples are obtained from publicly available repositories. dtangle works well even when these reference data sets were created using a different profiling technology. This points to scRNA-seq data as as a promising source for references.

dtangle has some of the same limitations as other algorithms. Primarily, it is necessary that the cell types comprising each sample be known in advance and that reference data is available. We will continue to develop dtangle to overcome some of these challenges to broaden its utility.

## Supplement

### Assessing The Relationship Between Actual and Measured Expression

One of the main components of dtangle’s approach is a linear model relating actual transcript concentration and measured gene expression. To explore it’s plausibility, we consider this linear model’s application to Affymetrix DNA microarray data and Illumina RNA-seq data.

#### Microarray Data

To explore the relationship between the amount of transcripts and the measured expression from microarray technology we consider the Latin Square data set from Affymetrix (Irizarry *et al.*, 2003). This data set was created by hybridizing a solution of complex human background mRNA with 42 transcripts spiked in at concentrations ranging from 0.125pM to 512pM. The spike-ins were done with 3 technical replicates of 14 hybridization experiments in a Latin square design. This data set lets us explore the relationship between measured expression and abundance of the transcripts because for each of the spiked-in transcripts we know both the expression measured by the array and the amount in which the transcript was spiked in.

The expression as measured by the microarray is best explained by a logistic fit in the spike-in amount (Supplementary Figure S11a). However the linear fit that dtangle assumes does quite well. The logistic fit has a slightly smaller *R*^2^ than the linear fit however there are several reasons we choose to model the relationship between spike-in amount and measured expression as linear. Firstly, the linear model is much simpler than the logistic model and has almost as good of a fit. For the linear model *R*^2^ = 0.957 while for the logistic least squares fit we have that *R*^2^ = 0.992. Thus we gain relatively little for using the more complex model. Furthermore, the simplicity of the linear fit can also be thought of as a regularization of the logistic model. The non-linearity of the logistic curve means it is very unstable model for extremely high or low measured expressions. That is, its inverse is undefined at or beyond these points. If the logistic model is used to estimate the true gene transcriptional abundance from measured expression data then small changes in measured expression might correspond to large changes in predicted amount. Indeed, the logistic curve will fail completely for measured expressions above its maximum or below its minimum. The linear model can be thought of as a regularized model between these two quantities. It makes sure that a linear change in measured expression will only ever affect a linear change in amount. While there is probably a true non-linear relationship between the expression measured by microarrays and the amounts of transcripts in the samples, a linear fit does quite well at approximating this relationship and is a regularized model for the truth.

#### RNA-seq

Another reason we favor a linear modeling of the relationship between amount and measured expression is because it is not only reasonable for microarray technology but a reasonable model for RNA-seq. To explore how our model interacts with RNA-seq technology we consider data from the Sequencing Quality Control project (SEQC Consortium, 2015). These data are available on GEO with accession GSE47774. Here we look at RNA-seq analysis run on Ambion ERCC Spike-In Control Mix 1 using Illumina HiSeq technology. The ERCC spike-in control mix contains 92 transcripts spiked-in at known concentrations. Hence this data set allows us to look at the relationship between measured expression and amount because both are known.

For this data the measured expression values (log2 of the read count plus one) are well approximated by a linear relationship to the spike-in concentration amount (Supplementary Figure S11b). Unlike the previously discussed microarray data the RNA-seq data does not seem well approximated by a logistic fit. For a simple linear regression we find *R*^2^ = 0.955 and so a linear fit seems reasonable.

#### Estimating The Slope

dtangle requires estimating the technological sensitivity parameter *γ* by some estimator 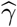. This *γ* is precisely the slope of the linear model we have just discussed (see Equation (3)). The value of 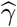 in dtangle’s algorithm may be set by the user if desired. However a pre-set value of 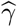 will be used by default if none is supplied. If 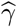 is not specified by the user, one need only specify the type of technology as either probe-level microarray, gene-level microarray, or RNA-seq. From here a default value of 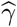 is chosen. These default values are estimated from spike-in experiments like those just discussed. The parameter *γ* is estimated separately on each of the different gene expression analysis technologies. For both RNA-seq and microarray spike-in data we fit regression models of measured expression on spike-in amount. These are the linear models seen in in the previous sections. We then take the median value of all the estimates of the slopes from each gene’s regression model. These form the estimate of 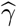. This is done for the RNA-seq data (on the log2 of the counts plus one) and microarray data (at the RMA-summarized gene level and raw log2 probe-level). These estimates set the default values for 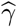 at .452 for probelevel microarrays, .699 for gene-level microarrays, and .943 for RNA-seq data. In any case dtangle seems robust to changes in 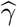 with best performance when 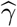 is between .5 and 1.

#### Slope Sensitivity

In order to evaluate the sensitivity of dtangle to changes in 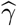 we conduct a meta-analysis of dtangle over many values for 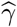 (Supplementary Figure S10). dtangle seems to perform poorly if 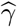 is less than about 1/2. However for 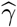 above about 1*/*2 dtangle is not particularly sensitive to the parameter.

---

**Figure S1:**
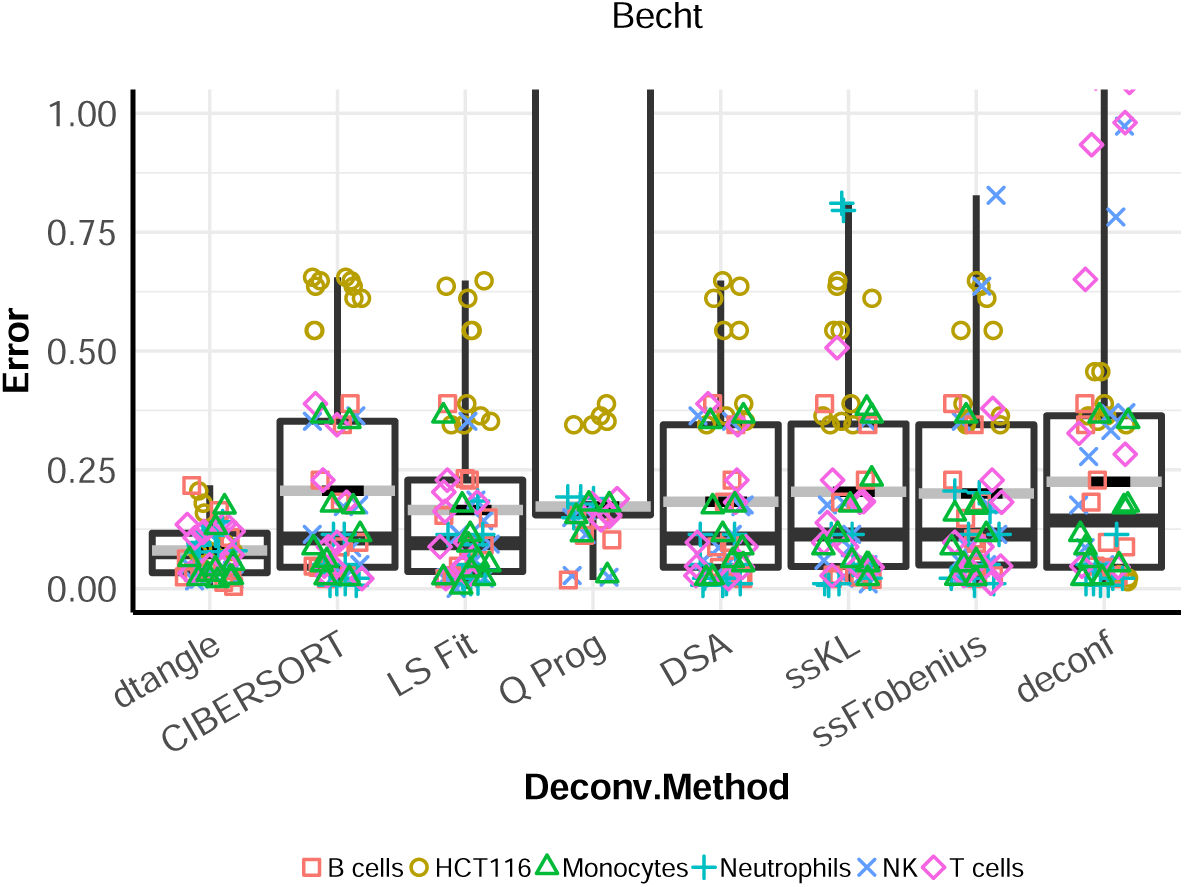
Comparing dtangle with other methods on Becht data. Side-by-side box plots of the absolute value of the errors for Becht data across each of the algorithms. The Q Prog. algorithm implementation fails for certain samples hence the very large errors.

**Table S1:** Eight deconvolution algorithms we compare.

| Deconvolution Methods |  |  |
| --- | --- | --- |
| Name | Citation |  |
| **dtangle** | (This publication) | } Partial |
| **CIBERSORT** | (Newman *et al.*, 2015) |  |
| **LS Fit** | (Abbas *et al.*, 2009) |  |
| **Q. Prog** | (Gong *et al.*, 2011) |  |
| **deconf** | (Repsilber *et al.*, 2010) | } Full |
| **DSA** | (Zhong *et al.*, 2013) |  |
| **ssFrobenius** | (Gaujoux and Seoighe, 2012) |  |
| **ssKL** | (Gaujoux and Seoighe, 2012) |  |

**Table S2:**
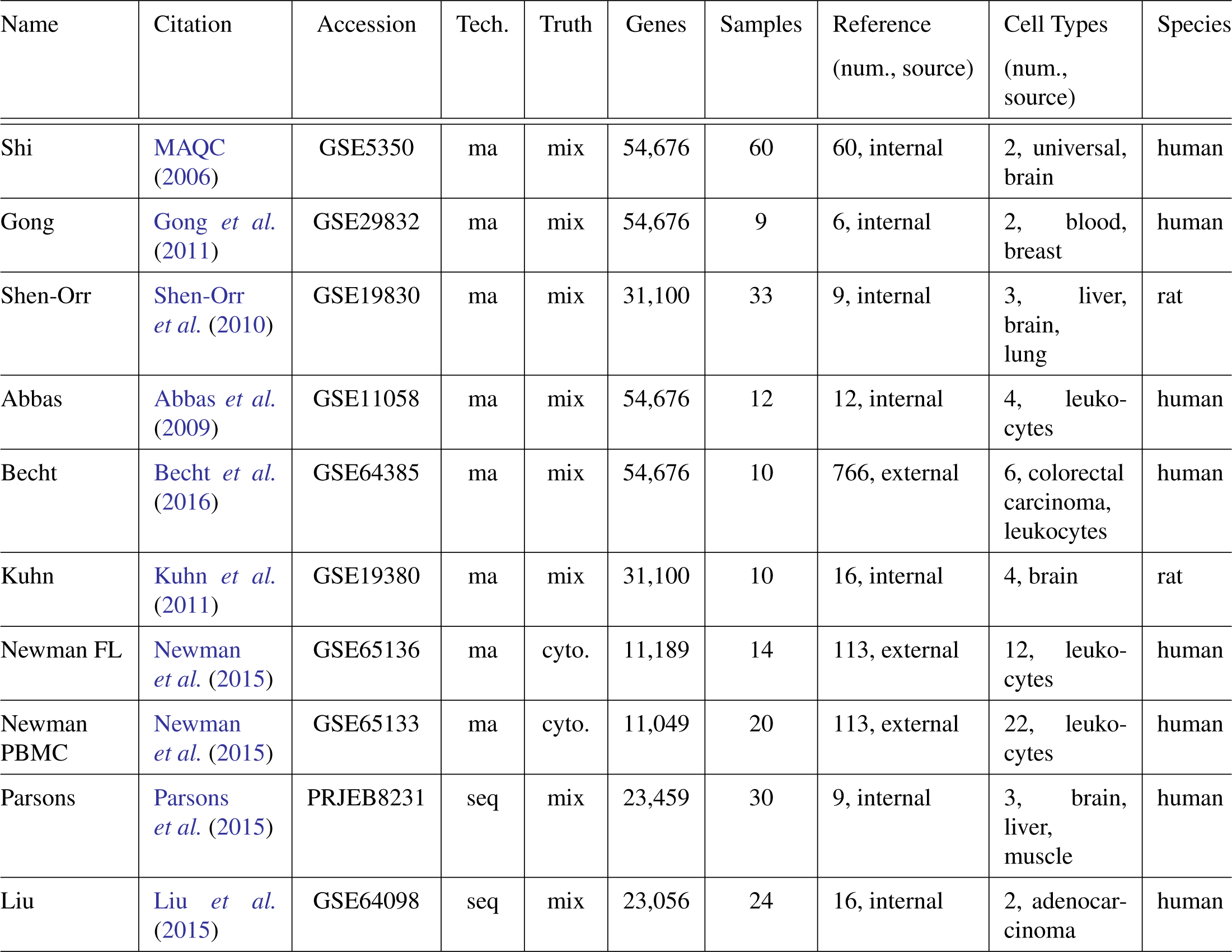
Benchmark data sets on which we compare deconvolution algorithms. The accession key is for GEO (or in the case of Parsons, ENA). The technology producing the data is either “ma” for microarray or “seq” for RNA-seq. The column “Truth” distinguishes between mixture experiments “mix” or data where the truth is known from flow cytometry “ctyo.” The number of gene expression measurements made by the technology is the column “Genes” and the number of unknown heterogeneous samples deconvolved is the column “Samples.” The column “Reference” lists the number of samples in the reference data along with the designation of “internal” if the pure reference samples were created part and parcel with the mixture experiment or “external” if the reference samples were collected from external data sources (typically GEO). The column “Cell Types” lists the number of cell types in the mixture samples and provides a description of the cell types along with the species from which the cell types come (in the column “Species”).

| Name | Citation | Accession | Tech. | Truth | Genes | Samples | Reference<br>(num., source) | Cell Types<br>(num., source) | Species |
| --- | --- | --- | --- | --- | --- | --- | --- | --- | --- |
| Shi | MAQC (2006) | GSE5350 | ma | mix | 54,676 | 60 | 60, internal | 2, universal, brain | human |
| Gong | Gong *et al.* (2011) | GSE29832 | ma | mix | 54,676 | 9 | 6, internal | 2, blood, breast | human |
| Shen-Orr | Shen-Orr *et al.* (2010) | GSE19830 | ma | mix | 31,100 | 33 | 9, internal | 3, liver, brain, lung | rat |
| Abbas | Abbas *et al.* (2009) | GSE11058 | ma | mix | 54,676 | 12 | 12, internal | 4, leukocytes | human |
| Becht | Becht *et al.* (2016) | GSE64385 | ma | mix | 54,676 | 10 | 766, external | 6, colorectal carcinoma, leukocytes | human |
| Kuhn | Kuhn *et al.* (2011) | GSE19380 | ma | mix | 31,100 | 10 | 16, internal | 4, brain | rat |
| Newman FL | Newman *et al.* (2015) | GSE65136 | ma | cyto. | 11,189 | 14 | 113, external | 12, leukocytes | human |
| Newman PBMC | Newman *et al.* (2015) | GSE65133 | ma | cyto. | 11,049 | 20 | 113, external | 22, leukocytes | human |
| Parsons | Parsons *et al.* (2015) | PRJEB8231 | seq | mix | 23,459 | 30 | 9, internal | 3, brain, liver, muscle | human |
| Liu | Liu *et al.* (2015) | GSE64098 | seq | mix | 23,056 | 24 | 16, internal | 2, adenocarcinoma | human |

**Figure S2:**
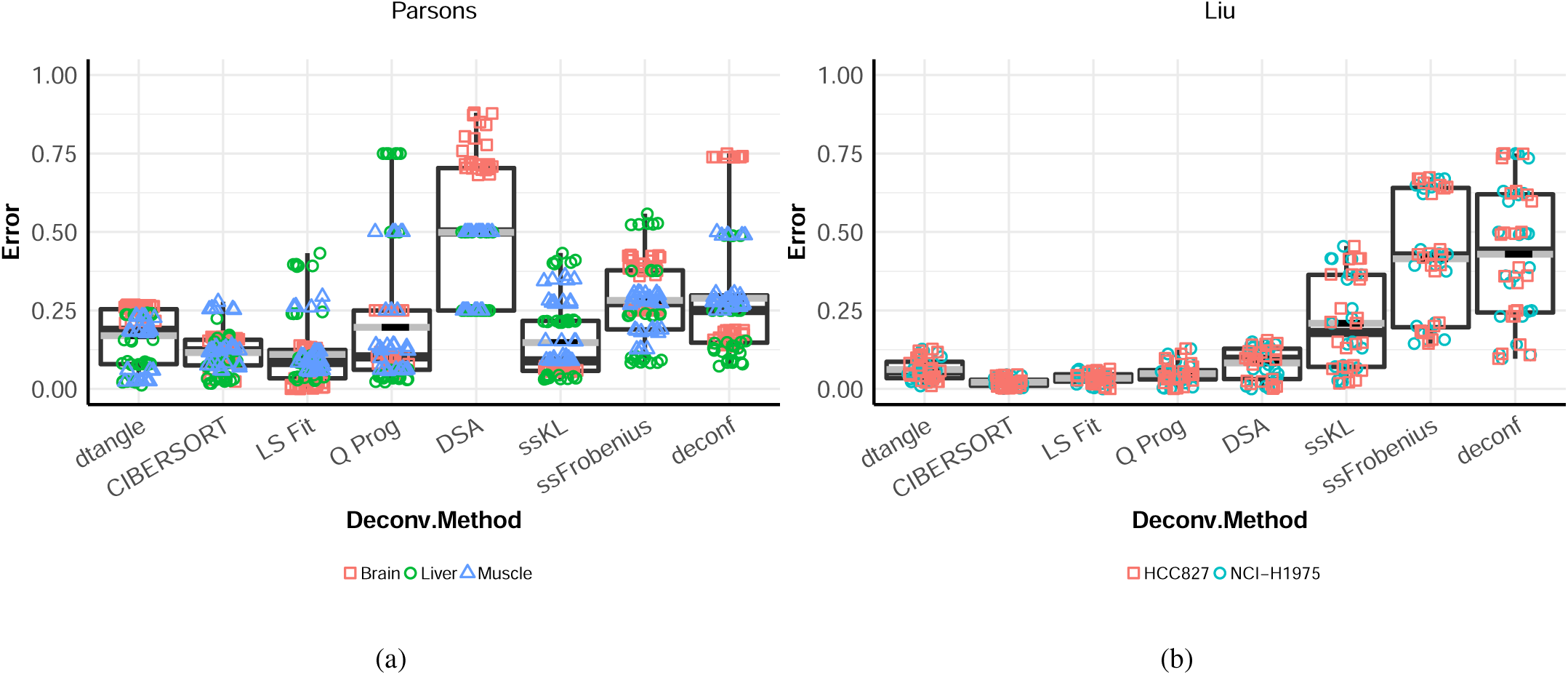
Side-by-side box plots of the absolute value of the errors for RNA-seq data sets Parsons (a) and Liu (b) across each of the algorithms.

**Figure S3:**
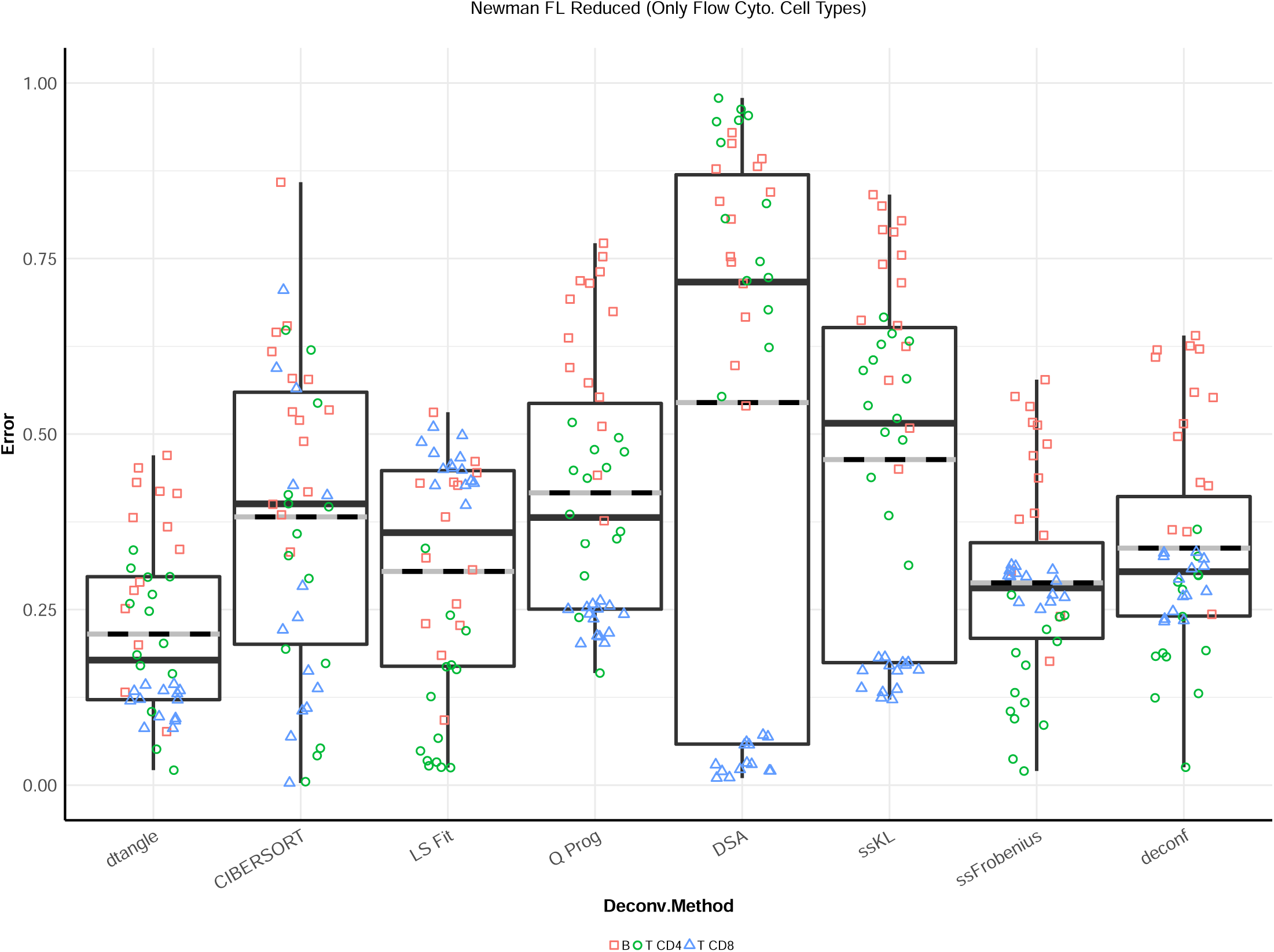
Side-by-side box plots of the absolute value of the errors for Newman flow cytometry data set Newman FL across each of the algorithms. This deconvolution has been done using only the three cell types that are reported by flow cytometry (B, CD4 T and CD8 T). The accuracy of the algorithms in predicting B cells types is higher than if we include cell types that are not reported by flow cytometry. Compare to Supplementary Figure S4.

**Figure S4:**
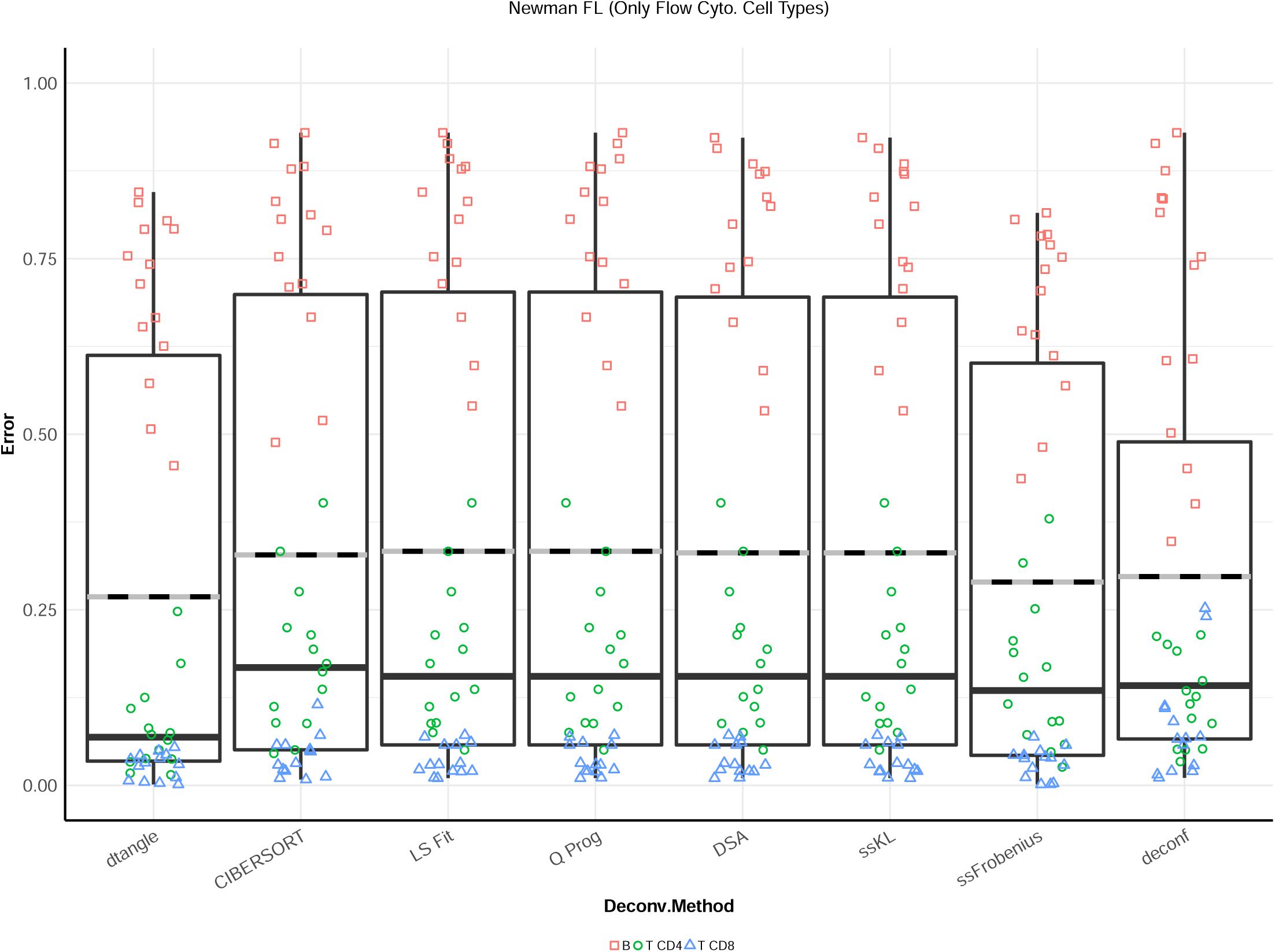
Side-by-side box plots of the absolute value of the errors for Newman flow cytometry data set Newman FL across each of the algorithms. This deconvolution has been done using 12 cell types (some of which are not reported by flow cytometry) however for the sake of comparison we only report the errors for the three cell types that are reported by flow cytometry (B, CD4 T and CD8 T). The inclusion in the deconvolution algorithms of cell types that are not reported by flow cytometry increases the error for the B cells. Compare to Supplementary Figure S3.

**Figure S5:**
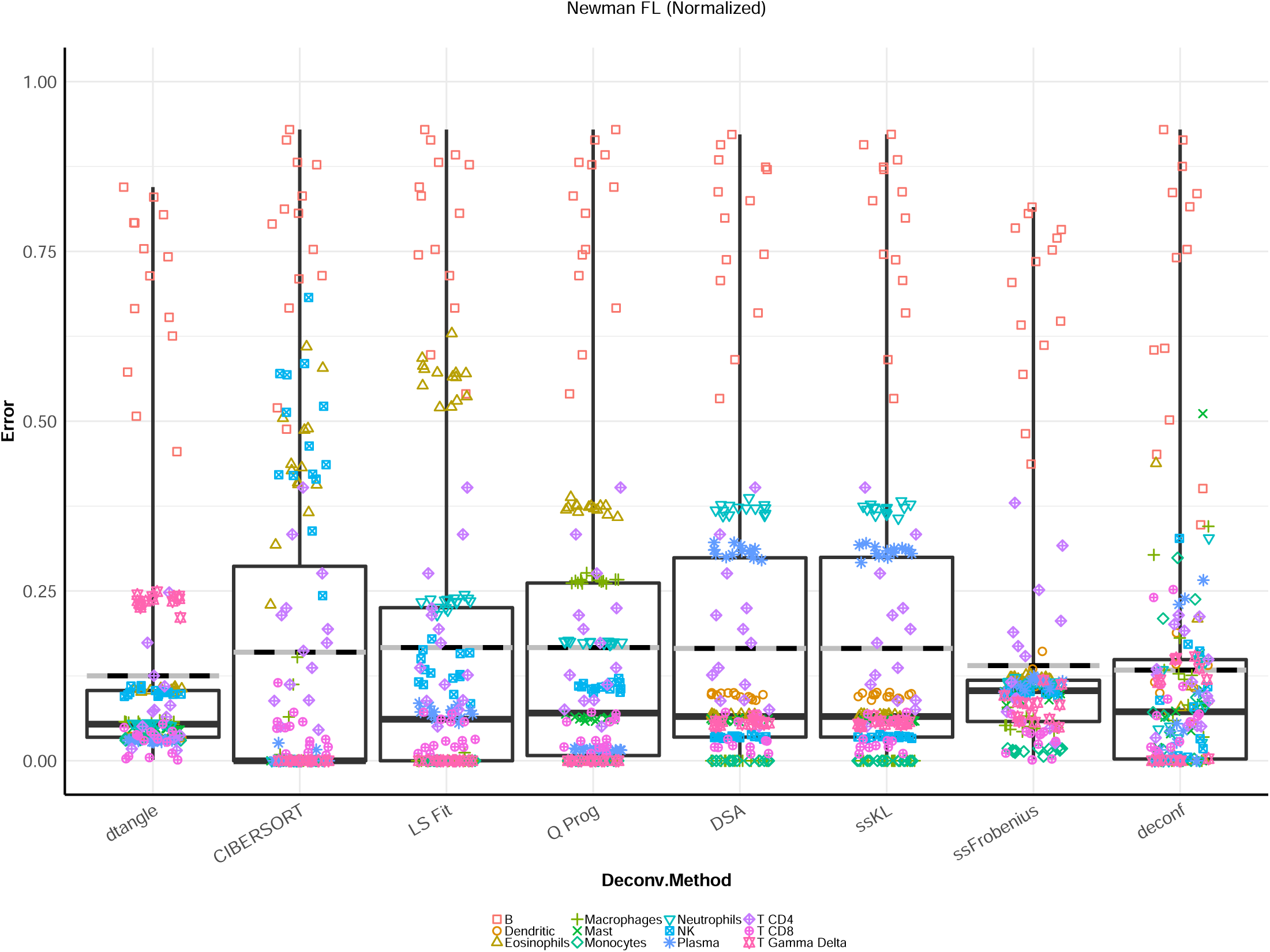
Side-by-side box plots of the absolute value of the errors for Newman flow cytometry data set Newman FL across each of the algorithms. Microarray data was quantile normalized before deconvolution. Compare to Supplementary Figure S6. While some cell types are predicted with more accuracy after quantile normalization the median error is increased by this normalization.

**Figure S6:**
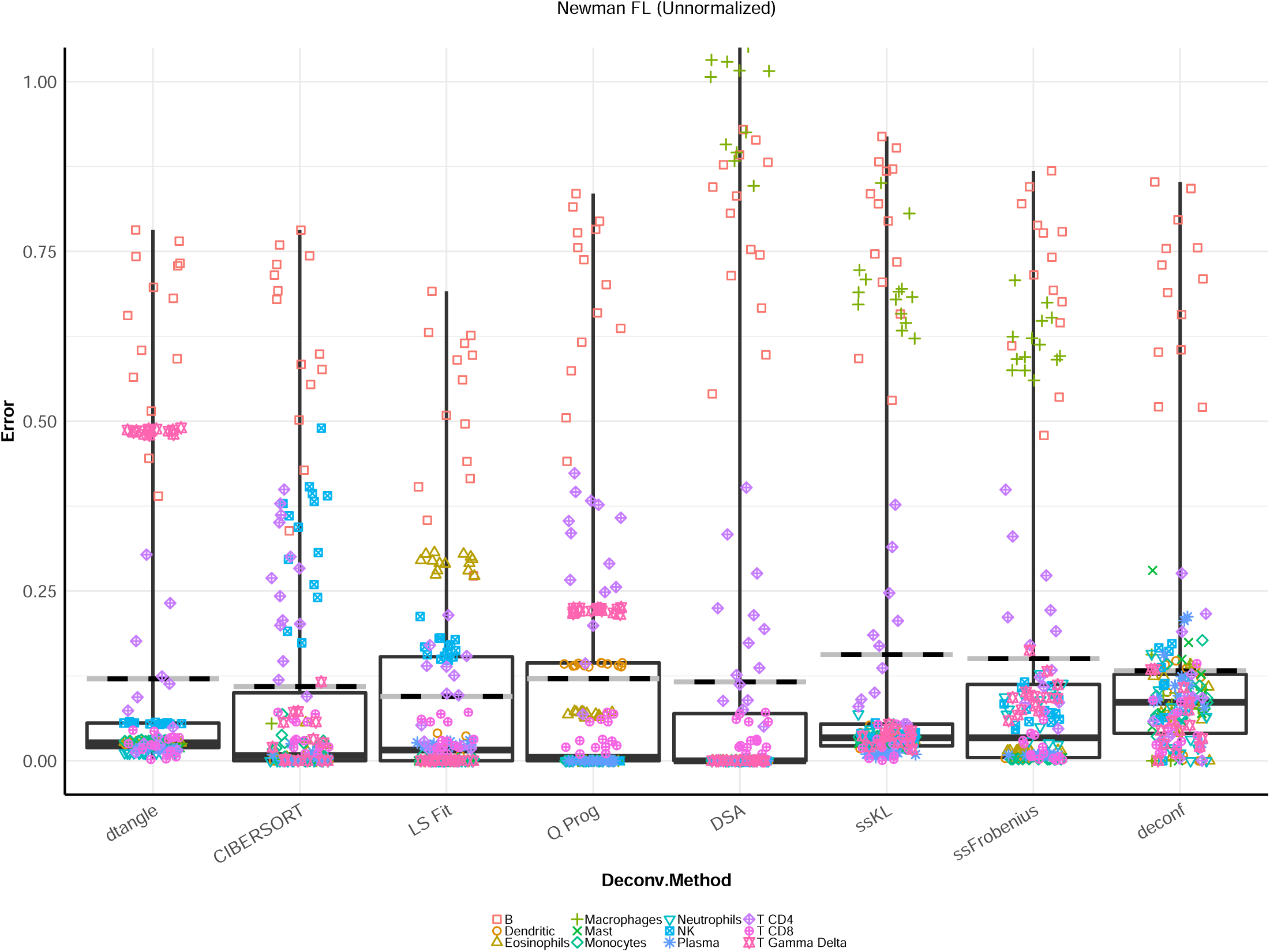
Side-by-side box plots of the absolute value of the errors for Newman flow cytometry data set Newman FL across each of the algorithms. Microarray data was **NOT** quantile normalized before deconvolution. Compare to Supplementary Figure S5. While the median error is lower without quantile normalization some cell types are predicted with significantly lower accuracy.

**Figure S7:**
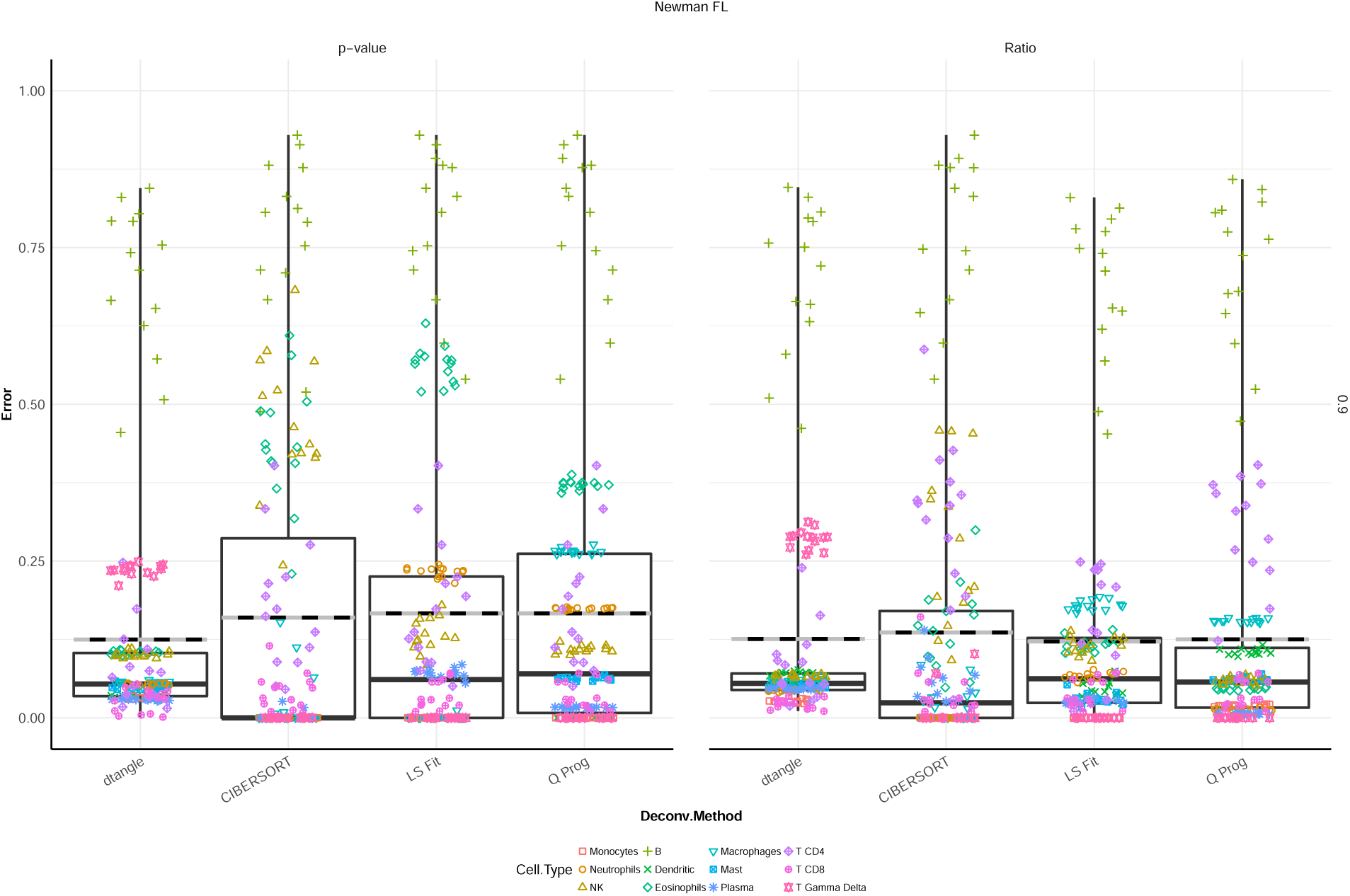
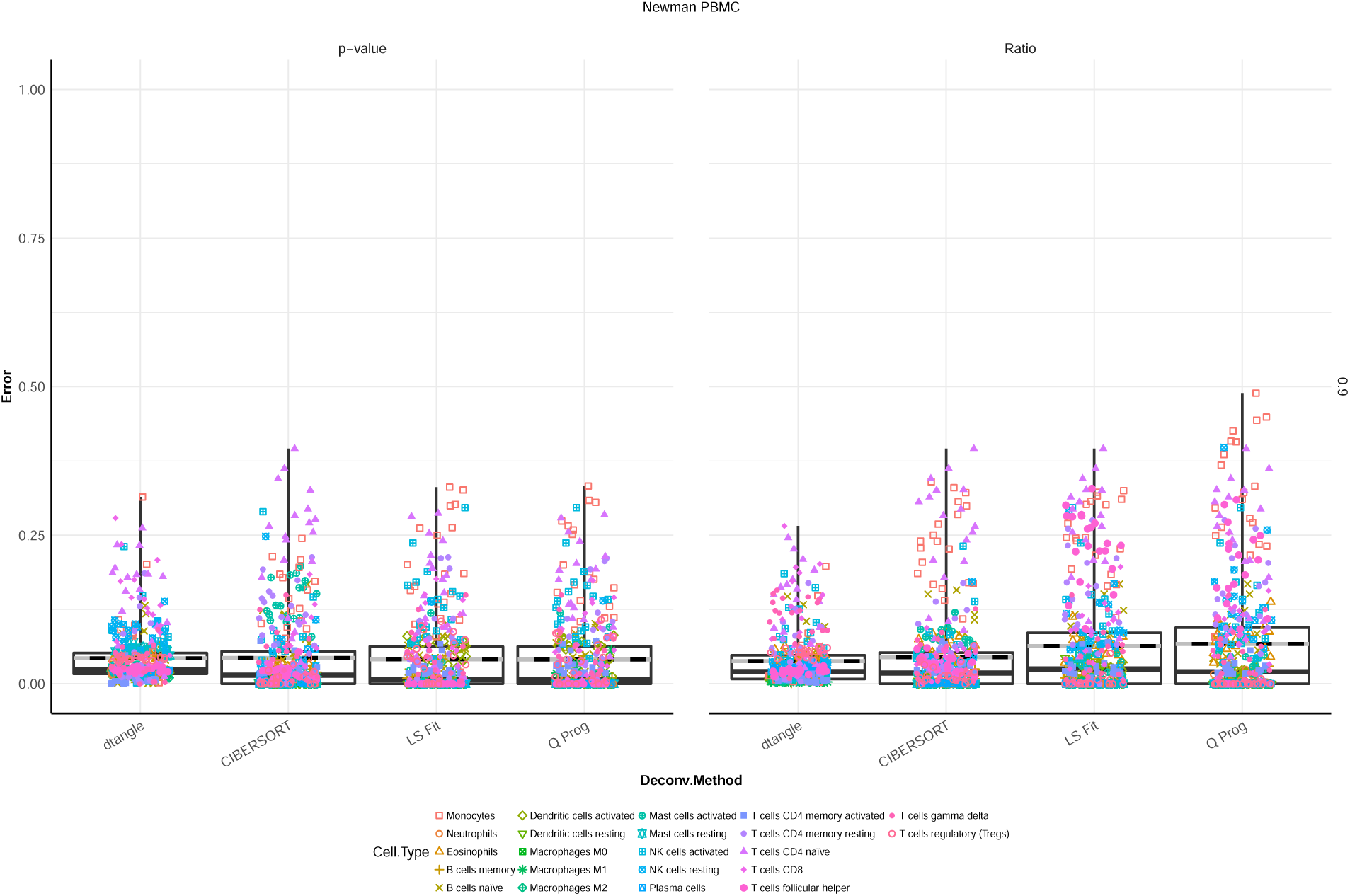
Full boxplots with legends for Newman data sets. (a) Boxplot of data from Newman et al. Slope for dtangle chosen automatically, markers chosen following Abbas et al. (p-value) and using the built-in method in dtangle (Ratio) using top 10% of identified markers in each case. (b) Boxplot of data from Newman et al. Slope for dtangle chosen automatically, markers chosen following Abbas et al. (p-value) and using the built-in method in dtangle (Ratio) using top 10% of identified markers in each case.

**Figure S8:**
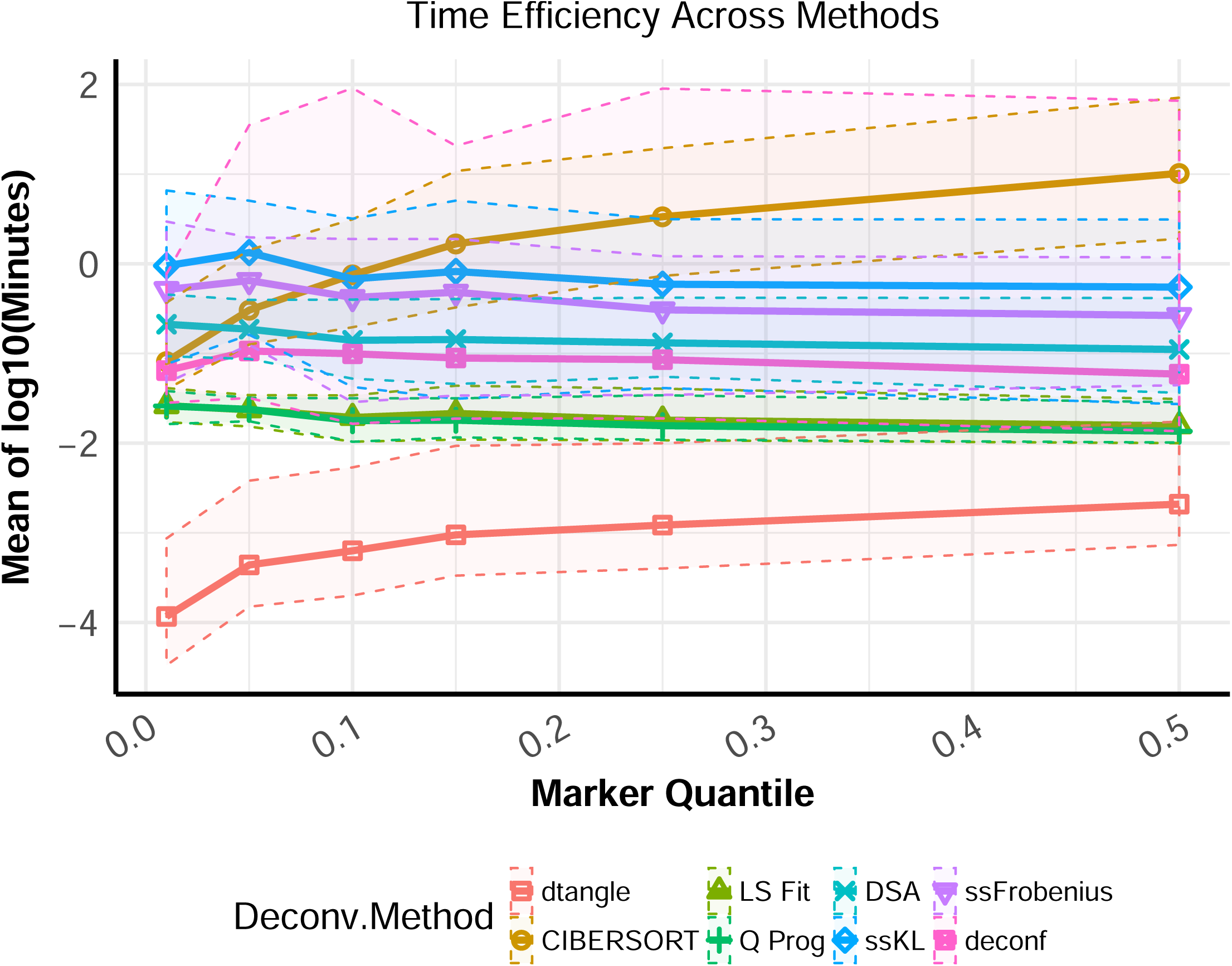
Mean of log_10_ of time (in minutes) each algorithm took to deconvolve all data sets. Maximum and minimum value error envelope is included.

**Figure S9:**
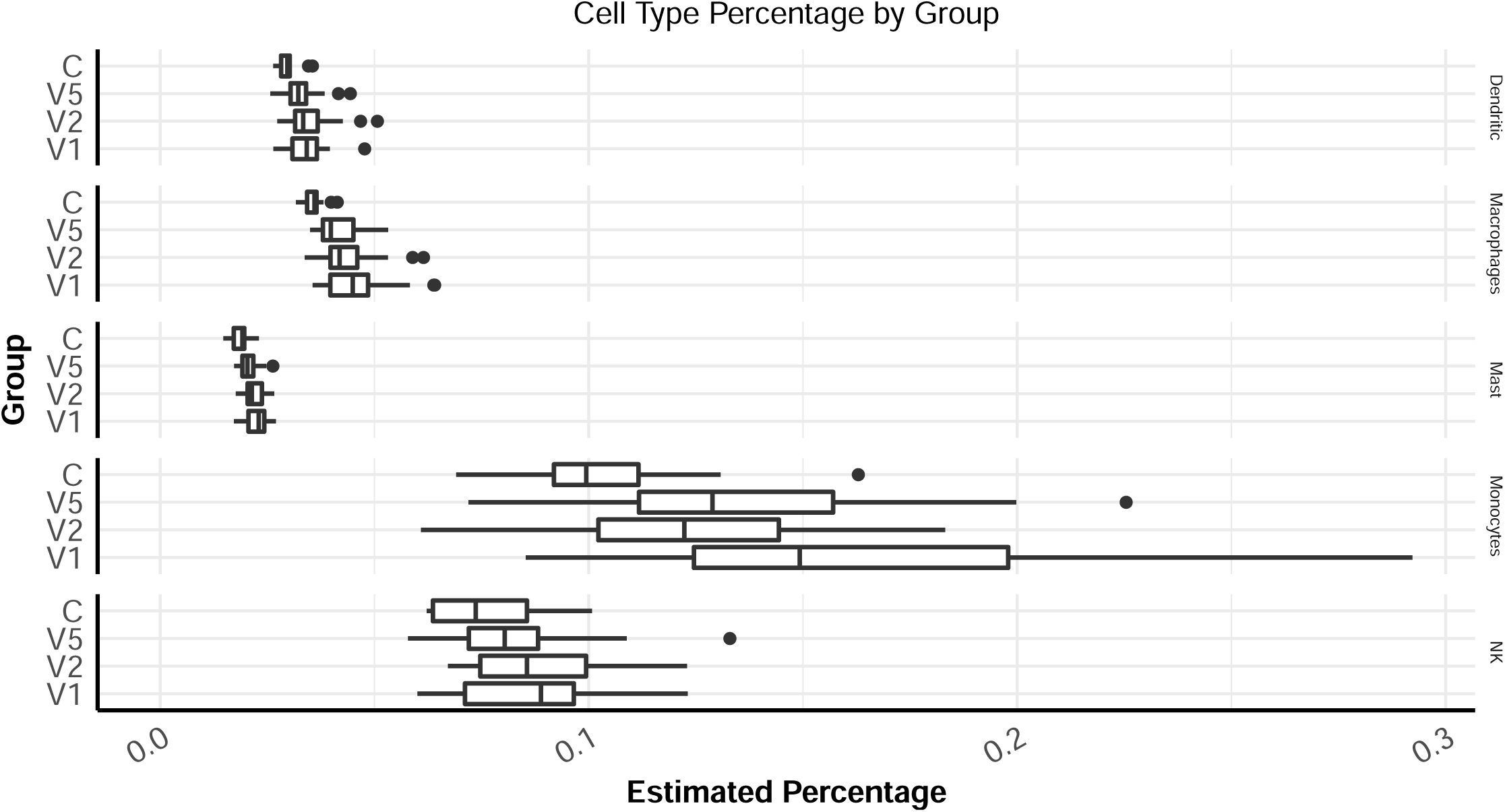
Estimated cell type proportions for the treatment group at times V1, V2 and V5 as well as the proportions for the control group. Plotted are estimated percentages for the professional phagocytes and the NK cells.

**Figure S10:**
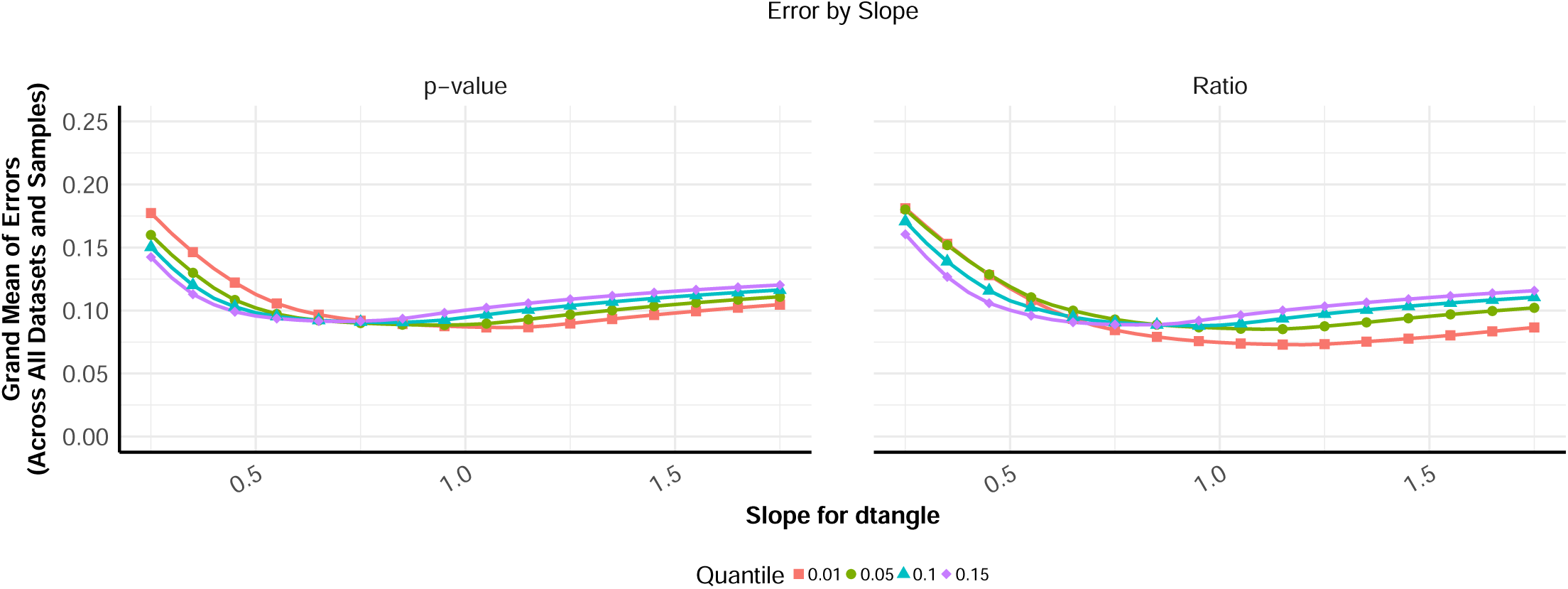
Meta-analysis slope for dtangle. Plotted is the grand error mean against the value of 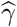 supplied to dtangle. This is done for several quantile cutoffs for markers (top 1%, 5%, 10% and 15%) as well as two methods for ranking marker genes (p-value and Ratio). dtangle is robust to choice of 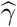 so long as it is above 1/2.

**Figure S11.**
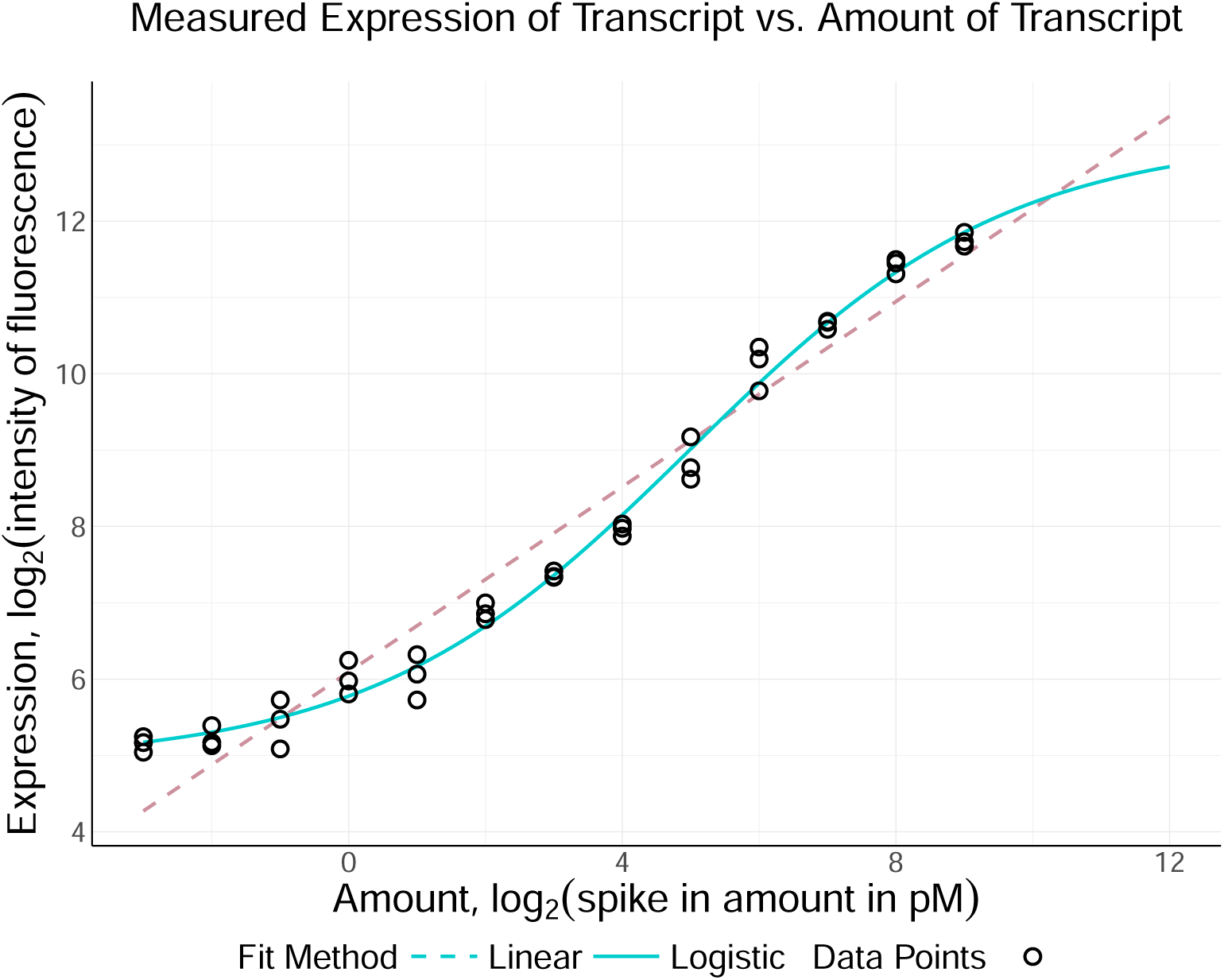
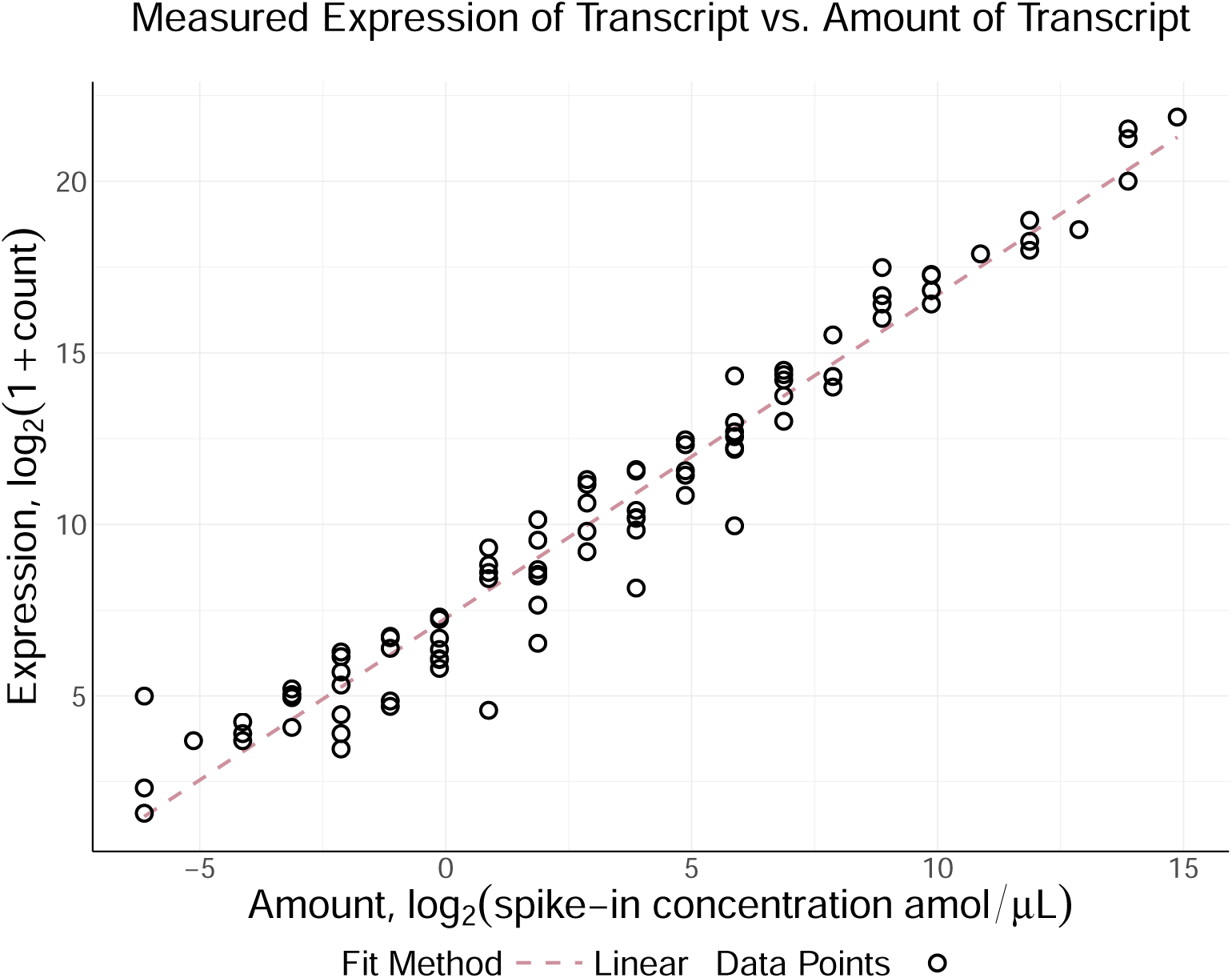
(a) log_2_ measured expression versus log_2_ concentration of a probe for gene TNFRSF1B. The relationship is approximated well by a linear model. While we have plotted amount against measured expression for one particular gene the results are generalizable to all genes. Points are plotted for the 13 experiments where the gene is spiked-in at a amount above zero and for each of the three technical replicates of each experiment. Along with the data points are plotted a linear and logistic least squares fit. The linear fit is a simple linear regression of measured expression on amount and the logistic fit is the least squares fit of a generalized logistic function of the form *β*_0_ + *β*_1_*/* (1 + exp (*β*_2_*x* + *β*_3_)). (b) log_2_ measured expression versus log_2_ concentration of ERCC spike-in controls in RNA-seq data. This relationship is highly linear. A linear least squares regression fit is plotted as a line.

